# Volume EM reveals three-dimensional architecture of the desmosome in epithelial cells and tissue models

**DOI:** 10.64898/2026.01.27.702185

**Authors:** Navaneetha Krishnan Bharathan, William Giang, Eunice Chen, Stephanie E. Zimmer, Sonam Lhamo, Danielle M. Jorgens, Jamie L. Inman, Vito Mennella, Manfred Auer, Andrew P. Kowalczyk

## Abstract

Desmosomes are a type of cell-cell adhesive junction present in cardiac tissue and epithelial tissues such as the epidermis. These intercellular junctions anchor to the intermediate filament cytoskeleton, providing mechanical integrity to the tissues in which they reside. Our understanding of desmosome architecture has largely been influenced by observations of two-dimensional images obtained through conventional electron microscopy. Here, using focused ion beam scanning electron microscopy, we report the three-dimensional ultrastructure of desmosomes in A431 and S1 human mammary epithelial cells. We also reveal differences in desmosome ultrastructure at homo- and heterotypic junctions of human nasal airway epithelial cells. Quantitative analyses of these volume EM datasets reveal variations in desmosome size, shape, and organization. Importantly, we report the presence of discontinuities or “holes” within the desmosome outer dense plaque, a novel feature that is observed in either one or both halves of a desmosome. This study provides the first comprehensive description of the epithelial desmosome as a three-dimensional structure, and emphasizes the need to investigate the effects of dynamic morphogenetic processes and disease states on desmosome ultrastructure.

## Introduction

First identified in the epidermis by light microscopy in the late 1800s, the desmosome has fascinated biologists studying the structures that connect neighboring cells (Bizzozero, 1864, 1870; Bharathan et al., 2024). The desmosome is mainly composed of proteins from three families: the cadherins, desmoglein and desmocollin; the armadillo proteins, plakoglobin (PG) and plakophilin (PKP); and the plakin protein desmoplakin (DP) (Bharathan et al., 2024; Zimmer and Kowalczyk, 2024). The cadherins engage in calcium-dependent interactions across the intercellular space and bind to PG and PKPs subjacent to the plasma membrane. PG and PKP both bind to DP, which in turn forms links with the keratin filament cytoskeleton to mechanically connect adjacent cells in a tissue. Defects in these proteins lead to a plethora of diseases affecting the skin and heart (Hegazy et al., 2022; Perl et al., 2024; Stahley and Kowalczyk, 2015), underscoring the need to understand desmosome formation and structure.

Electron microscopy (EM) has been used to study desmosome ultrastructure since the 1950s (Porter, 1956; Selby, 1955, 1957). In transmission electron micrographs, desmosomes appear as symmetrical structures consisting of five electron-dense stripes. This arrangement results in a “rail-road track” pattern of electron-dense stripes at the interface of two adjacent cells, with a total thickness of ∼100 nm and a diameter of 0.3–0.7 µm (Odland, 1958; North et al., 1999). The band subjacent to the plasma membrane is referred to as the outer dense plaque (ODP), while the more cytoplasmic band is called the inner dense plaque (IDP). Conventional TEM has been widely used to characterize variations in desmosome number, size/ diameter, and intercellular spacing between the plaques across stages of embryonic development (Overton, 1962; Bharathan and Dickinson, 2019). Further, immunogold EM and super-resolution imaging with dSTORM have revealed the organization of the major desmosomal proteins within the junction (North et al., 1999; Stahley et al., 2016; Beggs et al., 2022). In addition, electron tomography studies of mouse epidermal and liver desmosomes have provided angstrom-level resolution and suggest an ordered arrangement of the desmosomal cadherins (He et al., 2003; Sikora et al., 2020). However, these methods are hindered by inherent restraints such as low *z*-resolution due to thickness of serial sections and the tractability of imaging large samples, preventing one from appreciating the three-dimensional organization of the desmosome. Volume EM techniques, such as focused ion beam scanning electron microscopy (FIB-SEM), overcome these constraints by providing near-isotropic nanometer scale resolution of relatively larger samples. For example, FIB-SEM of the intercalated disc of murine cardiac ventricles revealed differences in desmosome size between wildtype and αT-catenin-deficient mice (Vanslembrouck et al., 2018, 2020). However, high-resolution three-dimensional (3-D) views of the epithelial desmosome are lacking.

In this study, using FIB-SEM datasets of A431 cells and additional epithelial cell culture models, we provide the first comprehensive, high-resolution, isotropic 3-D visualization and analysis of the epithelial desmosome. Reconstruction by segmentation and quantitative analysis reveal that desmosomes form in a variety of shapes and sizes, indicating a correlation between desmosome diameter and keratin filament association. We also observe differences in diameter and sphericity of the “typical” desmosome between various epithelial cell types. Finally, we report the presence of electron-lucent regions or gaps within the desmosome plaque. These findings provide a reference structure for normal desmosomes and a foundation to understand how these adhesive junctions are altered in disease states.

## Results

### Desmosome plaques display diversity in 3-D shape

Early EM studies described the desmosome as being ovoid in shape in en face views (Odland, 1958), but three-dimensional views of the epithelial desmosome are still lacking. We therefore sought to determine the 3-D architecture of the desmosome ODP using a structured illumination microscopy (Cryo-SIM) and focused ion beam scanning electron microscopy (FIB-SEM) workflow in A431 epithelial cells. To ensure near-native preservation of subcellular structures, samples were prepared by high-pressure freezing (HPF) (McDonald and Auer, 2006). To confirm desmosome regions, these cells stably express the desmosomal protein, desmoplakin, fluorescently-tagged to EGFP (DP-EGFP) (Godsel et al., 2005; Bharathan et al., 2023). With this workflow, we generated high-resolution volumetric views of the cell-cell interface, acquired at resolutions of either 4 nm or 8 nm isotropic voxels (Fig. 1a,b). In the high-resolution cryo-SIM images, desmoplakin (marked by DP-EGFP) exhibits a “railroad track” appearance at the cell-cell interface. Correlation of the light and electron microscopy (CLEM) images demonstrates that these “railroad track” desmoplakin that these “railroad track” desmoplakin puncta mark bona fide desmosomes (Fig. 1a).

**Figure 1.**
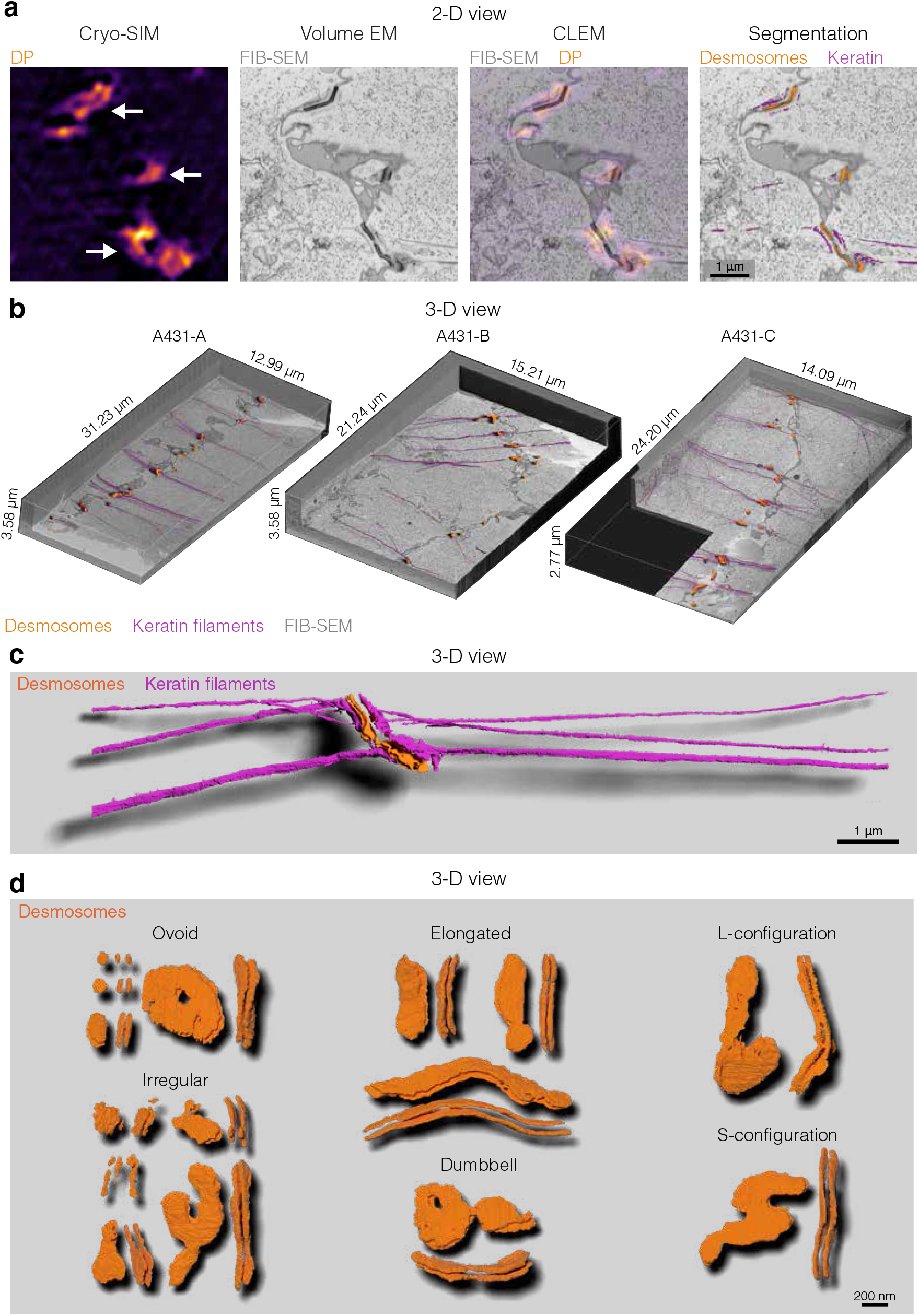
Desmosomes in A431 cells display various shapes and configurations. a) Alignment of DP-EGFP (desmoplakin) captured using cryo-SIM with the corresponding FIB-SEM orthoslice. b) 3-D views of the three A431 FIB-SEM datasets in this study. A431-A and A431-B were captured at 8 nm isotropic resolution. A431-C was captured at 4 nm isotropic resolution. Segmented desmosome outer dense plaques (orange) and keratin filaments (magenta) are also displayed. c) A three-dimensional rendering of the desmosome (orange) and the associated keratin filaments (magenta) on either side of the cell-cell contact. d) The various shapes of desmosome outer dense plaques found in the A431 datasets. The en face (where the full face of the ODP is in view) and a rotated view is shown for each desmosome.

Manual and semi-automated segmentation enabled us to efficiently label the electron-dense voxels that comprise the desmosome plaques. We then reconstructed these segmentations into 3-D objects to visualize the outer dense plaque (ODP) (Fig. 1c). Nearly all desmosomes possessed two ODPs, representing the two halves at the interface of adjacent cells (n=86/95). For some desmosomes, we were unable to reliably distinguish between individual ODP halves due to regions of electron density connecting the two halves (n=7/95). We next examined the three-dimensional architecture of each desmosome. In agreement with prior studies capturing en face views (Odland, 1958), most desmosomes are ovoid in shape. However, our data show that desmosomes also occur in elongated, “S”, or “L” configurations, or exhibit a “dumbbell” shape (Fig. 1d). A subset of desmosomes had more than two ODPs (n=2/95), where a plaque on one cell is paired with two plaques on another cell (Fig. 1d, “Irregular”). Interestingly, these desmosomes also appear larger in size than their “typical” ovoid counterparts, suggesting that such arrangements result from fusion of multiple smaller desmosomes. These high-resolution volumetric views reveal a previously unappreciated diversity in desmosome ultrastructure.

### Smaller ODPs typically lack intermediate filaments

We next performed quantitative analysis to obtain dimensions of the desmosome ODPs, including classical measurements such as the diameter and the distance between the two plaque halves. The maximum Feret diameter (MFD) indicates the longest distance between any two parallel tangents along each discrete object’s convex hull, which we used as a proxy for the ODP diameter. Combining data from all A431 datasets, we found that the diameter of individual ODPs ranged between 30-1850 nm, with a median diameter of ∼260 nm (n=187 ODPs in 95 desmosomes). We next segmented and examined the keratin filament network and their association with each half of the desmosome. As we have previously reported (Bharathan et al., 2023), some desmosomes possess anchored filaments in only one cell of the cell-cell pair, while others completely lacked any association with identifiable keratin filaments (Fig. 2a,b). Further analysis revealed that desmosomes possessing any keratin filaments (on one or both sides), likely indicating mature desmosomes, have a median diameter = 462 nm (n=44/95). In contrast, desmosomes that lacked any identifiable keratin filaments were relatively smaller (median diameter = 178 nm, n=51/95) (Fig. 2a-D).

**Figure 2.**
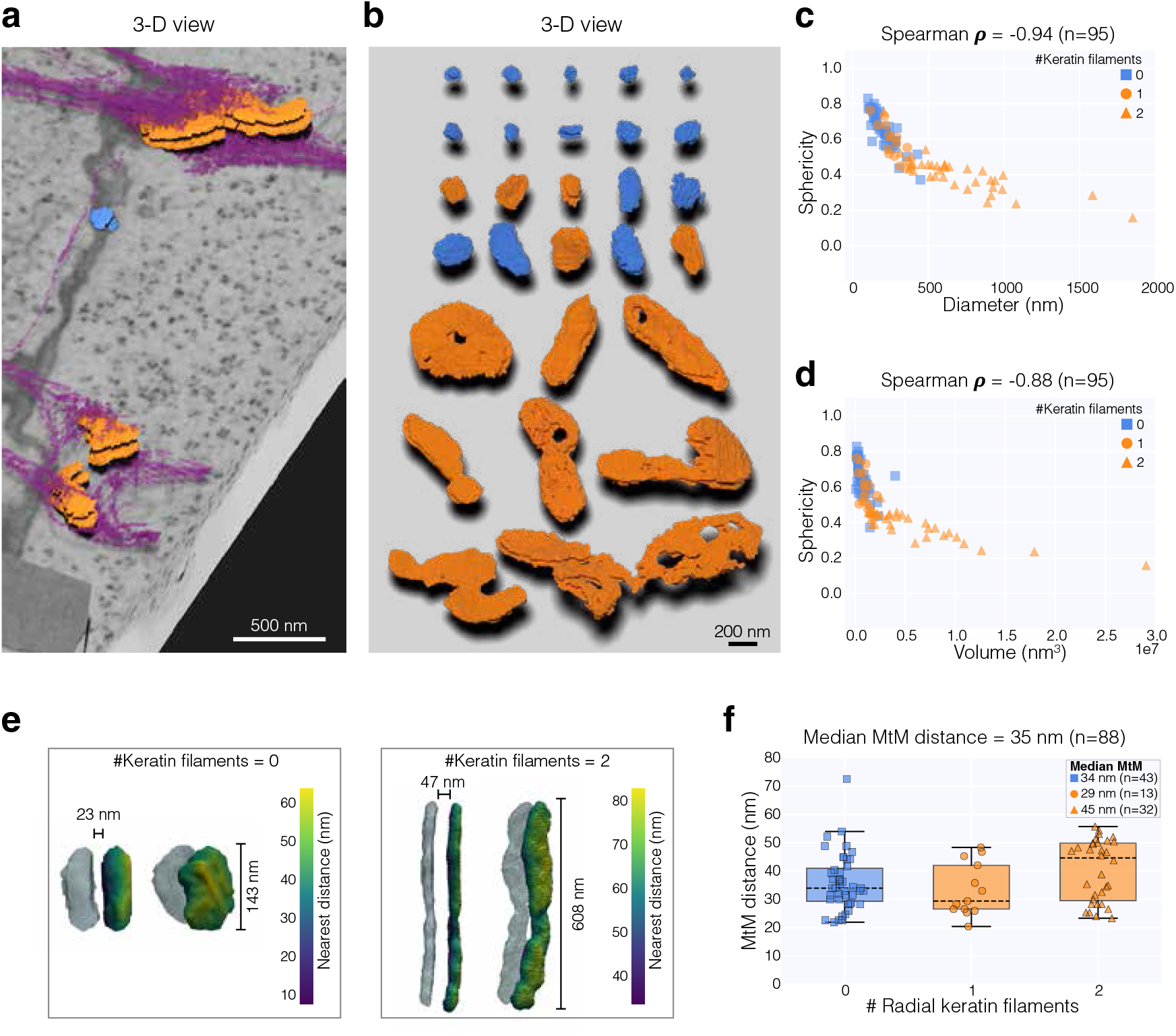
Desmosomes in A431 cells show differential association with keratin filaments. a) A 3-D view of desmosomes (orange) associating with keratin filaments (magenta) and desmosomes with no keratin filament association (blue) in a region of the A431 FIB-SEM dataset. b) Representative views of desmosomes color-coded to indicate association with keratin filaments (orange) or lack of association (blue). c-d) Scatter plots depicting sphericity vs. diameter (c) or volume (d). Data points are color-coded by desmosomes having no keratin filaments (blue square), having keratin filaments on only one side (orange circle), or having keratin filaments on either side (orange triangle). Correlation coefficients are also shown. e) Examples of membrane-to-membrane (MtM) distance measurements of two desmosomes. Color bar denotes the nearest neighbor distance of a vertex point in one plaque with a vertex point in the paired plaque. Corresponding en face view is shown on the right. f) Box-and-whisker plots depicting the MtM distances of various desmosomes in the A431 datasets. Dashed line represents the median, and upper and lower bounds represent the 75th and 25th percentiles, respectively. Whiskers extend from the box to the farthest data point lying within 1.5x of the inter-quartile range. Points outside the whiskers are statistical outliers.

We next estimated the sphericity (Φ) of the individual ODPs as a proxy for its aspect ratio. Sphericity is the degree to which a 3-D object approaches a sphere and is measured by calculating the ratio of the surface area of a sphere with the same volume as the object being measured to the actual surface area of the object. A perfect sphere has a Φ = 1, while objects that are elongated or having an irregular shape have a Φ value closer to 0. In A431 cells, the sphericity of individual ODP plaques ranged between 0.159-0.95 (n=187 ODPs in 95 desmosomes), reflecting the diversity in ODP shapes we observe (Fig. 1d, 2b). We next wanted to evaluate whether there was a correlation between ODP diameter (i.e., MFD) and sphericity. As indicated in Fig. 2c, there is a robust inverse relationship between these two variables. Similarly, there is a robust inverse relationship between sphericity and volume (Fig. 2d). Altogether, these findings suggest that longer, larger desmosomes tend to have less sphericity, while shorter, smaller desmosomes tend to approach a spherical shape.

We also determined the width of the extracellular space between both plaques of a whole desmosome, i.e., the “membrane-to-membrane” (MtM) distance. The median MtM distance was estimated to be ∼35 nm (n=88) (Fig. 2e). These measurements fall within MtM distances obtained via electron tomography (ET) and molecular dynamics simulations of Cryo-ET data (He et al., 2003; Al-Amoudi et al., 2007; Sikora et al., 2020). Interestingly, we found that desmosomes with keratin filaments on either side displayed larger median MtM distances relative to desmosomes with keratin filaments on only one side or those lacking filaments (Fig. 2f).

### Desmosomes exhibit “holes” or “tunnels” within regions of the plaque

Further inspection of the 3-D reconstructed desmosome revealed gaps within the ODP, where only the plasma membrane is observed. In some cases, these electron-lucent plasma membrane gaps are only present in one plaque which we term “holes” (Fig. 3a). In other instances, the plaques can have symmetrically-aligned gaps across both halves. We term these arrangements “‘tunnels”. To better visualize the concept of ODP holes and tunnels, we made use of an imaginary light source to cast a shadow of either the whole desmosome (all plaques) or of its individual plaques (Fig. 3a). Tunnels coincide with electron-lucent regions located laterally between two electron-dense regions that would otherwise suggest the presence of two separate desmosomes (Fig. 3b). We categorized ODPs into 4 groups: 1) no holes/tunnels; 2) only holes (in either one or both ODPs); 3) only tunnels; 4) both holes and tunnels. We next wanted to determine whether there was a relationship between ODP structural characteristics and the presence of holes and tunnels. Scatter plots comparing sphericity to diameter or volume revealed that smaller desmosomes tend to lack any holes/tunnels, while larger desmosomes have tunnels and/or holes (Fig. 3c,d). These features might thus represent an intermediate step during fission or during fusion of multiple smaller desmosomes.

**Figure 3.**
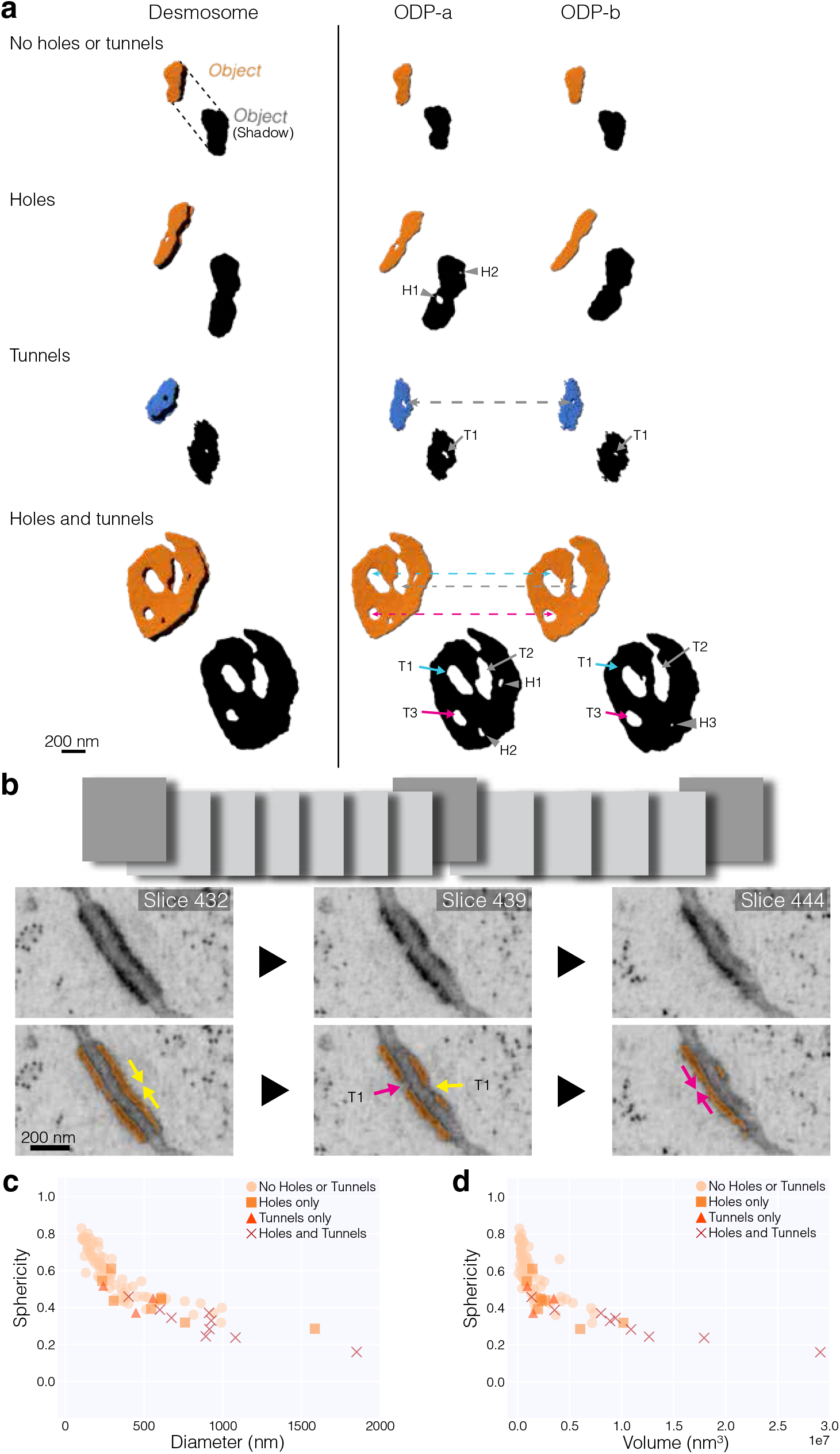
Desmosomes in A431 cells have holes and tunnels. a) Desmosomes in A431 cells have no holes, holes in only one plaque, holes in both plaques forming a tunnel, or holes and tunnels. The holes and tunnels within the whole desmosome (left) and each plaque of a desmosome (right) are visualized with a projection of its shadow. b) Representative FIB-SEM orthoslices of a desmosome (orange). A single slice shows a gap between adjacent densities on both cells appearing as two adjacent desmosomes (slice 439, middle). Viewing slices above or below this slice clearly show that the electron densities within each cell connect (paired yellow arrows or pink arrows). The gap (middle panel) is a hole within each plaque, forming a tunnel. c-d) Scatter plots depicting sphericity vs. diameter (c) or volume (d). Data points are color-coded by desmosomes having no holes or tunnels (circles), having holes only (squares), having only tunnels (triangles), or having both holes and tunnels (red crosses).

### FIB-SEM reveals morphological differences between desmosomes of airway and S1 acini epithelia

Having characterized the size, shape, and organization of desmosomes in the A431 epithelial cell model, we next wanted to determine desmosome ultrastructural characteristics in other cell/tissue types acquired through volume EM. We first segmented desmosome ODPs in acinar structures formed from a non-malignant human mammary epithelial cell (HMEC) line, HMT-3522-S1 (S1), in two FIB-SEM datasets, designated “HMEC-A” (Jorgens et al., 2017) and “HMEC-B”. While both datasets had a large number of desmosomes, we only segmented complete desmosomes, i.e., those in which we could identify the top and bottom boundaries, resulting in 23 ODPs. All desmosomes in these acini have keratin filament attachment on both halves (Fig. 4a). Similar to desmosomes in A431 cells, these S1 acini desmosomes display a variety of shapes and sizes, with ODPs of many large desmosomes displaying holes and tunnels (Fig. 4b). In both S1 acini datasets, desmosomes are larger than those found in A431 cells, with median diameters of 549 nm (HMEC-A) and 374 nm (HMEC-B), with some desmosomes exceeding 1950 nm in diameter. The median Φ was 0.38 (HMEC-A) and 0.43 (HMEC-B), indicating elongated or irregular organization. Correlative analyses again revealed that sphericity shared a robust inverse relationship with both ODP diameter and volume (Fig. 4c,d). Interestingly, we found that the median MtM distance (∼44 nm) was larger than those in A431 cells (Fig. 4e). These data indicate that there are cell-type specific differences in desmosome architecture. However, it is possible that this variation might be driven in part by differences in fixation method between the A431 and HMEC datasets.

**Figure 4.**
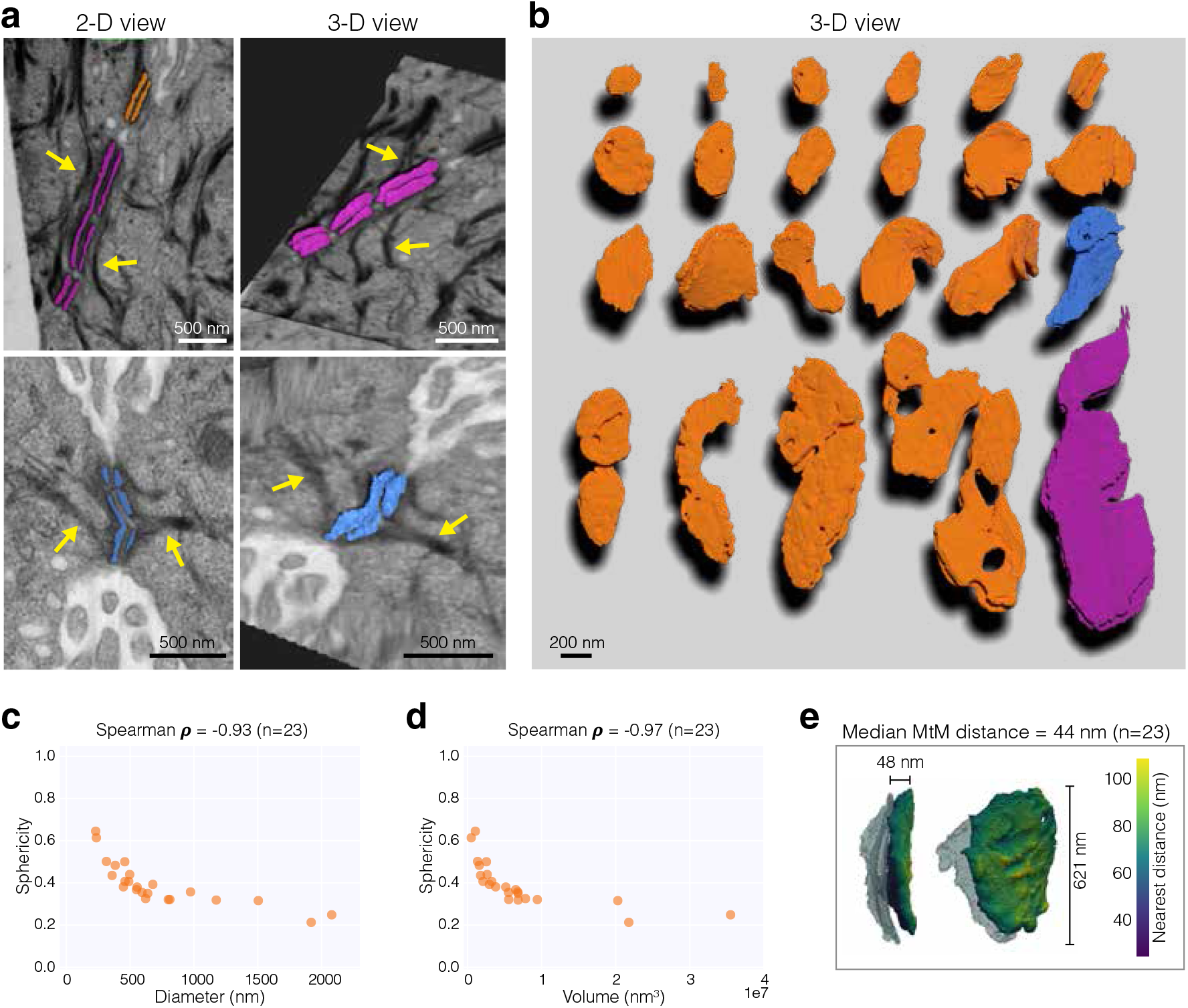
Desmosomes in HMEC S1 acini. a) Representative 2-D (left) and 3-D (right) views of segmented desmosomes (orange, magenta, blue) in FIB-SEM datasets of HMEC S1 acini. Yellow arrows point to keratin filaments. b) 3-D views of all 23 desmosomes in the dataset. The whole desmosome including all constituent plaques are shown. Blue and magenta desmosomes are the same objects depicted in (a). c-d) Scatter plots depicting sphericity vs. diameter (c) or volume (d) of HMEC desmosomes (orange). Correlation coefficients are also shown. e) An example of membrane-to-membrane (MtM) distance measurements of a desmosome in HMEC acini. Color bar denotes the nearest neighbor distance of a vertex point in one plaque with a vertex point in the paired plaque. Corresponding en face view is shown on the right.

Lastly, we segmented desmosomes in human airway pseudostratified epithelium, a multicellular tissue lining the respiratory tract which play a critical function in removing bacteria, viruses and pollutants through mucociliary clearance. The human airway pseudostratified epithelium analyzed was differentiated from primary nasal basal stem cell progenitors Air Liquid Interface (ALI) cell culture (Vijayakumaran et al., 2024). Prior studies have demonstrated that nasal airway epithelial cells share similar organization to the bronchial region and can be used surrogates for the pseudostratified epithelium of the respiratory tract in vivo (Devalia et al., 1990; McDougall et al., 2008). This dataset consists mainly of fully differentiated multiciliated cells (MCC), which are involved in mucociliary clearance in airway epithelia (Liu et al., 2020; Davis and Wypych, 2021), and contain early differentiating/deuterosomal (DC) and mucus producing goblet cells (GC). Additionally, this dataset contains a cell exhibiting morphological similarities to tuft cells (TC), although its identity could not be conclusively determined. All cell types have prominent tight junctions (TJ) at the apical regions of cell-cell contact, with a belt of adherens junctions (AJ) immediately basal to the TJs (Fig. 5a). We segmented these junctions along with desmosomes, which are located basal to both AJs and TJs on the lateral cell-cell contacts (Fig. 5b). We also segregated and analyzed desmosomes present between homotypic (MCC–MCC) or heterotypic (MCC–GC, MCC– DC, MCC–TC, DC–TC) cell-cell interfaces (n=209 desmosomes) (Fig. 5b-d). We found that these airway epithelia desmosomes exhibit a median diameter of ∼124 nm, much smaller than desmosomes of A431 and S1 acini. There were minimal differences in the range of diameters between different cell-cell interaction types. In contrast to the other epithelial cell datasets, we found no holes or tunnels in airway epithelia desmosomes. In addition, keratin filaments were observed but could not be reliably segmented. The correlation coefficients between sphericity (median Ф o= 0.86) and both ODP size and volume were weaker in this dataset (Fig. 5e). Further, airway epithelial desmosomes exhibit a median MtM distance of ∼58 nm (Fig. 5f,g), larger than those observed in A431 cells and S1 acini. We also plotted the desmosome diameter against relative *z*-position along the lateral cell-cell borders. We restricted this analysis to cell-cell borders between an MCC–MCC pair and an MCC–DC pair (Fig. 6a,b). Interestingly, the desmosomes at both borders were larger at the most apical region relative to the ones below (Fig. 6c,d). In addition, there appeared to be a somewhat robust correlation between desmosome size and relative *z*-position among desmosomes at the interface of the MCC–GC cell pair (Fig. 6d). Together, these results suggest that desmosome ODPs generally approach a spherical shape within cells of airway epithelia. In addition, our data reveal for the first time the heterogeneity of desmosomes in multiple cell types in the human airway pseudostratified epithelium.

**Figure 5.**
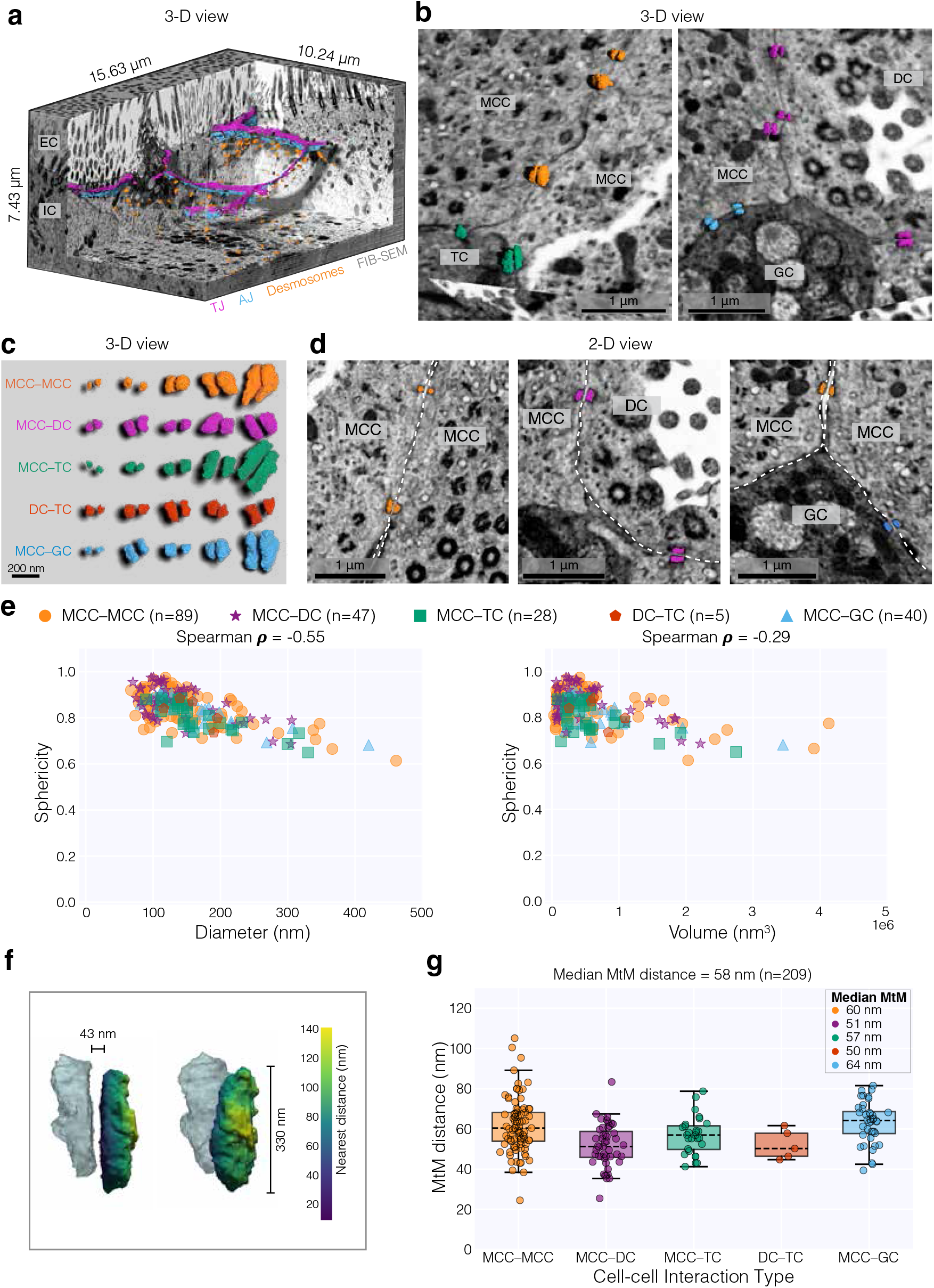
Desmosomes in human nasal basal airway epithelia. a) 3-D views of the airway epithelia FIB-SEM dataset captured at 10 nm isotropic resolution. Segmented desmosome outer dense plaques (orange), adherens junctions (blue), and tight junctions (magenta) are also displayed. EC: Extracellular space; IC: Intracellular regions. b-d) Representative 3-D views (b,c) or 2-D orthoslices (d) of segmented desmosomes between cell-cell pairs in airway epithelia. Desmosomes are color-coded by specific cell-cell pair. MCC: Multiciliated cell; DC: Deuterosomal cell; TC: Tuft cell; GC: Goblet cell. Dashed white lines depict the plasma membrane. e) Scatter plots depicting sphericity vs. mean Feret diameter (left) or volume (right) of airway epithelial desmosomes. Data points are color-coded by cell-cell pair. Correlation coefficients are also shown. f) An example of membrane-to-membrane (MtM) distance measurements of a desmosome in airway epithelia. Color bar denotes the nearest neighbor distance of a vertex point in one plaque with a vertex point in the paired plaque. Corresponding en face view is shown on the right. g) Box-and-whisker plots depicting the MtM distances of various desmosomes in the airway epithelia datasets. Dashed line represents the median, and upper and lower bounds represent the 75th and 25th percentiles, respectively. Whiskers extend from the box to the farthest data point lying within 1.5x of the inter-quartile range. Points outside the whiskers are statistical outliers.

**Figure 6.**
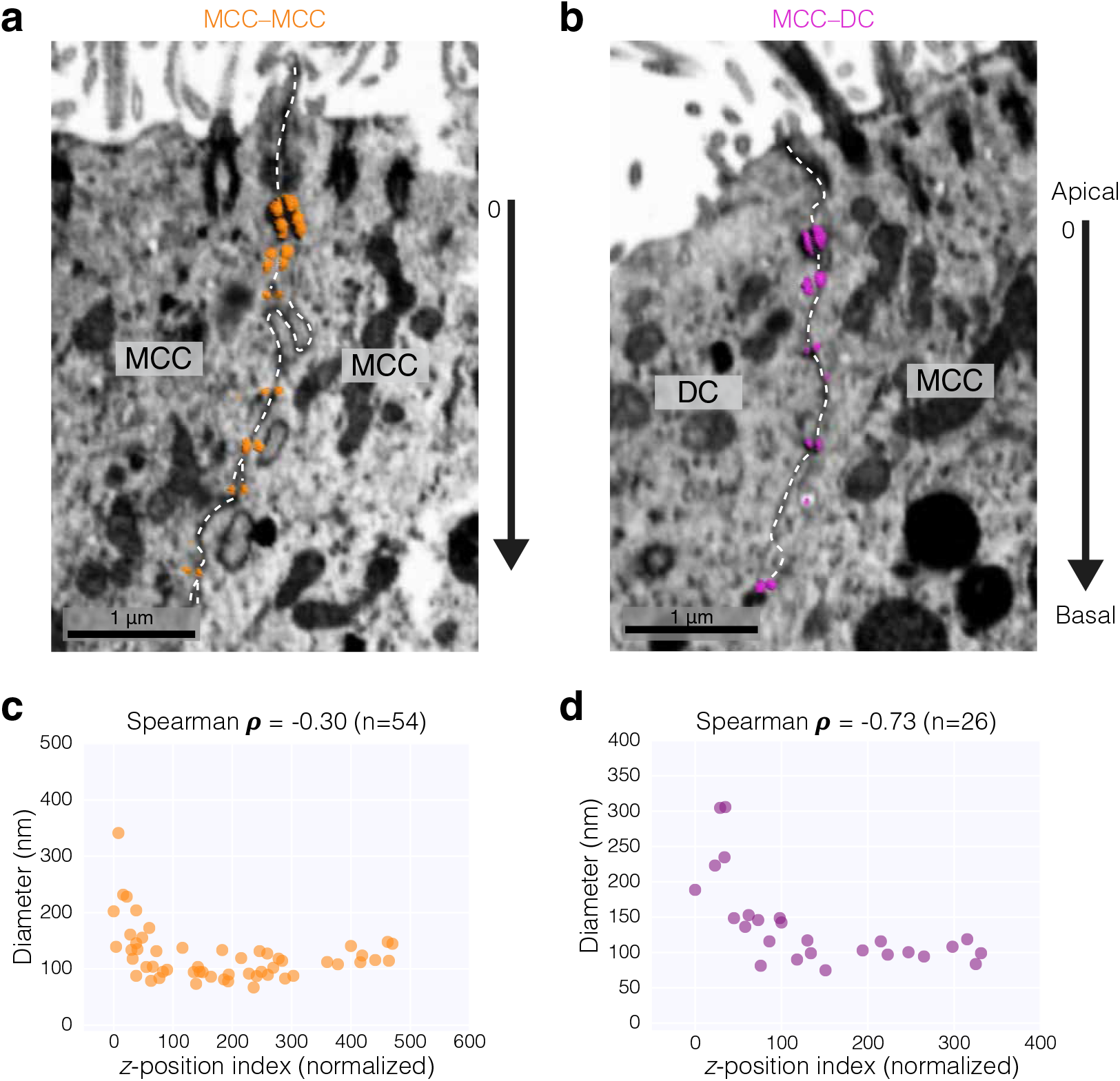
Desmosome size varies along the lateral cell-cell border. a-b) Representative images of FIB-SEM data and segmented desmosomes between an MCC–MCC pair (orange, a) or a DC–MCC pair (magenta, b). The z-index of the most apical desmosome is set as the reference point and assigned a value of 0. The *z*-indices of desmosomes basal to the most apical desmosome are normalized to 0. Dashed white lines depict the plasma membrane. c-d) Scatter plots depicting mean Feret diameter vs. *z*-position index of each desmosome between the MCC–MCC pair (a) and the DC–MCC pair (b). Data points are color-coded by cell-cell pair. Correlation coefficients are also shown.

## Discussion

We employed FIB-SEM to achieve high-resolution volumetric views of the epithelial desmosome derived from different tissues. These datasets reveal the diversity and complexity in configurations of the desmosome across cell types. Quantitative analysis of reconstructed 3-D segmentations show that these junctional complexes exhibit a range of sizes, with the largest desmosomes having a diameter over 10X higher in length relative to the smallest ones. Further, while small desmosomes tend to be ovoid, larger desmosomes have irregular shapes. On comparing multiple datasets, our analyses reveal that, unlike cells in airway epithelia, A431 epidermoid carcinoma cell monolayers and HMEC acini can form desmosomes greater than 500 nm. Similarly, the distance between adjacent plaques varies greatly between cell types. Importantly, we describe for the first time the presence of holes within the desmosome plaque in both A431 monolayers and HMEC acini.

The near-isotropic and high resolution of the volumes used in this study allowed for the detection of electron-lucent regions within electron-dense desmosome plaques, being observed in one or both plaque halves. These holes and tunnels are prevalent among the larger desmosomes in our datasets and are absent in smaller desmosomes. Notably, this phenomenon does not appear to be an artifact of fixation or cell type, since these structures are found in two distinct datasets collected by different groups using either 2-D or 3-D cell culture models and prepared using different fixation methods. Though this work is the first report of holes or tunnels within desmosomes, Kelly and Shienvold (Kelly and Shienvold, 1976) documented ‘discontinuities of particular mat’ in freeze-fracture images from newt epidermis which have a similar appearance to the structures described here. We have since found similar, unnoted discontinuities in other published freeze-fracture EM images from various organisms and tissue samples as well (Breathnach, 1975; Breathnach et al., 1976; Elias and Friend, 1976; Wolburg et al., 1983; Skepper, 1989), suggesting that these holes and tunnels may be a long-overlooked phenomenon.

Classical EM studies acquiring images of a single slice of desmosomes might have captured tunnels, but these regions were likely interpreted as gaps between distinct desmosomes. Resolution-limited light microscopy techniques might have also missed these structures as they are smaller than the diffraction limit. Our finding that desmosome holes and tunnels exist in a range of sizes leads us to speculate that these structures might have functional significance. Desmosomes are highly stable structures but also engage in dynamic behaviors, including lateral movements along the membrane and fusion of individual desmosomes into larger assemblies (Windoffer et al., 2002; Gloushankova et al., 2003; Bharathan et al., 2023). Desmosomal complexes are also protein-dense lipid raft domains in which non-desmosomal proteins are likely excluded based on specific physicochemical properties (Stahley et al., 2014; Lewis et al., 2019; Zimmer and Kowalczyk, 2020, 2024). It is tempting to imagine these holes and tunnels might be conduits for cell-cell communication, mechanisms for tension relief, or mechano-signaling platforms. However, given the heterogenous environment of the plasma membrane, desmosome holes and tunnels may simply be randomly isolated regions of the membrane created by desmosome fusion events.

As discussed, desmosomes anchor to the underlying intermediate filament network in a pairwise manner on either side of the junction, facilitated by the cytoplasmic desmosomal protein, desmoplakin (DP). Our data show that larger desmosomes tend to be associated with thick keratin filament bundles, while most desmosomes <300 nm in diameter display no detectable filaments. Live-cell imaging of HaCaT keratinocytes has shown that nascent desmosomes serve as nucleation sites for the assembly and elongation of keratin filaments, which increase in stability as desmosomes mature (Moch et al., 2020). However, the limited resolution inherent to light microscopy might preclude detection of keratin filament anchorage to nascent or immature desmosomes. These observations, taken together with our data presented here suggest a sequence of events where desmosome protein assembly occurs first, followed by recruitment and attachment of keratin filaments as desmosomal protein clustering increases. The smaller desmosomes in the current study might reflect nascent desmosome assemblies that have not yet been anchored to keratin filaments.

We segmented desmosomes in airway pseudostratified epithelium differentiated from human primary nasal basal stem cell progenitors (Vijayakumaran et al., 2024). Importantly, we provide high-resolution volumetric views of the desmosome at heterotypic cell-cell junctions, i.e., between goblet cells and multiciliated cells. Although desmosome organization appeared largely similar between homotypic and heterotypic cell-cell interfaces, differences might become apparent in pulmonary disease. Notably, *DSP* (desmoplakin) was identified as a genetic risk locus in a genome-wide association study of idiopathic pulmonary fibrosis patients (Fingerlin et al., 2013), and subsequent studies have revealed multiple *DSP* alleles with altered expression associated with an increased risk of lung disease (Mathai et al., 2016). Loss of DP in bronchial epithelial cells was also found to result in increased cell migration and increased expression of genes associated with the ECM and EMT (Hao et al., 2020). In addition, Dsp loss in *Xenopus laevis* embryos driven by antisense morpholinos resulted in defective radial intercalation between epidermal layers (Bharathan and Dickinson, 2019). This morphogenetic process is also thought to occur in mammalian airway epithelia (Rock et al., 2009). These studies underscore the need to elucidate desmosome ultrastructure during the development and differentiation of airway epithelia.

Overall, these findings reveal the complexity of desmosome ultrastructure that would have been difficult to visualize with conventional 2-D EM techniques. We expect these insights to stimulate studies examining whether and how desmosome protein composition, tissue-type, calcium dependence, mechanical stress, developmental stage, and disease impact desmosome 3-D organization.

## Materials and Methods

### Volume EM imaging of A431 epithelial cells

In this manuscript, we process previously-acquired FIB-SEM imaging data of a monolayer culture of A431 cells. A protocol detailing the culturing, preparation, imaging, and processing of these cells was published previously (Bharathan et al., 2023). Briefly, A431 cells expressing DP-EGFP and mApple-VAPB were cultured on sapphire disks (3mm diameter, 50-80 μm thick; Nanjing Co-Energy Optical Crystal Co., Ltd.) and stained with MitoView 650 (70075; Biotium; working conc. 25-500 nM) in a 37°C/5% CO_2_ incubator for 15 min and washed. Samples were cryofixed using High Pressure Freezing (HPF Compact 01; Wohlwend GmbH). Samples were then stored in liquid N_2_. Epifluorescence images were taken to identify regions of interest (ROIs), followed by imaging using 3-D structured illumination microscopy (3-D-SIM). Samples were then stained with 2% OsO_4_, 0.1% Uranyl Acetate, and 3% water in acetone under liquid N_2_, freeze substituted, and resin embedded. A micro X-ray CT system (XRadia 510, Carl Zeiss X-ray Microscopy, Inc.) was used to identify the positions of the cells of interest. Samples were sputter coated with gold and carbon, and then imaged on a Zeiss Gemini 450 Field Emission SEM and a Zeiss Capella FIB column oriented at 90 degrees to the SEM beam (Xu et al., 2017). Images were acquired at either 8 nm or 4 nm isotropic resolution. FIB-SEM images were then aligned and reconstructed followed by CLEM registration to align fluorescence and EM image data.

### FIB-SEM of S1 HMEC acini

One dataset was previously published (designated as “HMEC-A”), and another dataset has not been previously published (designated as “HMEC-B”). Images were acquired at either 4 nm isotropic resolution (HMEC-A) or at 5nm x 5nm x 10nm (HMEC-B) resolution as described previously (Jorgens et al., 2017). The actual voxel size for the HMEC-A dataset was 3.7nm x 4.9nm x 4nm. The HMEC-A dataset had errors during acquisition (detailed further in (Jorgens et al., 2017)) and were thus chopped into two tiles along the *z-*axis. Regions which had sample tearing or other artefacts were not processed further. To further reduce file sizes, one of the two HMEC-A tiles was chopped into two smaller tiles. All tiles in both HMEC-A and HMEC-B datasets were then drift-corrected in Fiji using the Correct 3D Drift plugin (Parslow et al., 2014). Briefly, image stacks were loaded into Fiji, the scale was calibrated, and the hyperstack was re-ordered to switch the slices (*z*) and frames (*t*). Images in the stack were drift-corrected by specifying the Max shift at either 100 or 250 pixels (for *x, y*, and *z*). Finally, the hyperstack was re-ordered to switch the slices and frames back to the original setting.

### FIB-SEM of nasal airway epithelial cells

Images were acquired at 10 nm isotropic resolution as described previously (Vijayakumaran et al., 2024), and were downloaded from OpenOrganelle at the highest resolution. The dataset was further cropped to reduce file size; remove regions with sample tearing, imaging artefacts, or resin; and to remove slices without cell-cell contacts.

### FIB-SEM image processing

Datasets were then resliced (rotated) to an orientation akin to typical light microscopy z-stacks. To reduce file sizes, the dataset acquired at 4 nm isotropic resolution (“A431-C”) was divided into twelve tiles of equal dimensions, one of which was previously published (Bharathan et al., 2023). Similarly, the dataset acquired at 8 nm isotropic resolution was cropped to include two tiles with cell-cell borders (“A431-A” and “A431-B”).

Keratin intermediate filaments (only for A431 datasets) and the desmosome outer dense plaque were segmented using the pixel classification workflow from ilastik (v1.4.1) or using a 3D U-Net as described previously (Lhamo et al., 2025). Segmentations were cleaned up using Microscopy Image Browser v2.9103 (MIB) as previously described (Belevich et al., 2016; Bharathan et al., 2023). Each desmosome (consisting of its component plaques on either side of the junction) was assigned as a separate object. Segmentation results were exported from MIB as 3-D TIFs and imported into Dragonfly 3D World software (Ver. 2025.1 for Windows) along with the corresponding cropped FIB-SEM dataset.

Three-dimensional renderings were generated using Blender 4.5 and Dragonfly 3D World software (Comet Technologies Canada Inc. (2025). Dragonfly 3D World., 2025).

### FIB-SEM image analysis

In Dragonfly 3D World, all desmosome objects in a dataset were merged into a “Multi-ROI”, with each desmosome having its own index value. Connected component analysis (26-connection) was then done on this “Multi-ROI” to generate separate objects, i.e., individual plaques. The following measurements of the individual objects were computed in Dragonfly: Maximum Feret Diameter (MFD), Volume, Sphericity. These measurements were then cross-indexed to the “Multi-ROI” to match measured values of individual plaques to those of the corresponding pair. The statistics were exported as comma separated value files. For correlation statistics, individual plaques were treated as separate desmosomes and corresponding values were compared. To represent the length of a whole desmosome as a single measurement, the MFD value of the largest ODP was used. Graphs were plotted using seaborn (v0.13.2) and matplotlib (3.10.0) (Waskom, 2021; Hunter, 2007; Anaconda Software Distribution.).

To measure membrane-to-membrane (MtM) distances, contour meshes for all desmosome objects were generated with a Threshold = 50 and exported using macros in Dragonfly 3D World. A k-dimensional tree was used for fast nearest-neighbor calculations between the surfaces of the two plaques. For reducing the MtM distance of 3-D surfaces into a single measurement, the lower quartile of the distance distribution was determined to be a robust metric in the “A431-C” dataset.

To compare the *z*-position and diameters of the cell-cell borders in the airway epithelia dataset, the actual *z*-coordinates of each desmosome, i.e., “Center of Mass Index Z”, were obtained with Dragonfly 3D World. These values were then normalized such that the most apical desmosome had a *z*-position index of 0, using the formula:

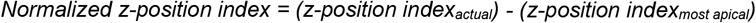

### Statistics and Reproducibility

Some desmosomes in the FIB-SEM datasets were excluded because of sample tearing or because only part of the desmosome was contained within the cropped datasets. Further cropping of FIB-SEM datasets was necessary for analysis due to computer hardware constraints and segmentation requirements. The cropped regions for analysis were chosen because they contained desmosomes without being greatly impacted by sample tearing.

Statistical testing was performed using custom Python scripting (v.3.10.14) and executed in a Jupyter Notebook environment. Relationships between variables were assessed using Spearman’s rank correlation coefficient (*ρ*). Python scripts will be made available on an online repository.

## Data availability

All code and macros will be made available on an online repository. A431 and airway epithelia FIB-SEM datasets are hosted on:

https://www.openorganelle.org/datasets/aic_desmosome-2,

https://www.openorganelle.org/datasets/aic_desmosome-3,

https://openorganelle.janelia.org/datasets/cam_hum-airway-14500.

Other imaging source data will be made available on an online repository. All other data supporting the findings of this study are available from the corresponding author on reasonable request.

## Acknowledgements

This work was supported by: National Institutes of Health grant R01AR048266 (A.P.K.); Children’s Miracle Network Faculty Research grant 1110000016 (N.K.B.). Cryo-SIM and FIB-SEM imaging of A431 cells were done in collaboration with the Advanced Imaging Center at Janelia Research Campus, a facility jointly supported by the Gordon and Betty Moore Foundation and the Howard Hughes Medical Institute. HMEC-A and HMEC-B FIB-SEM data collection was performed at the Oregon Health and Science University Multiscale Microscopy Core.

## References

1. Al-Amoudi, A., D.C. Díez, M.J. Betts, and A.S. Frangakis. 2007. The molecular architecture of cadherins in native epidermal desmosomes. Nature. 450:832–837. doi:10.1038/NATURE05994.

2. Anaconda Software Distribution.

3. Beggs, R.R., T.C. Rao, W.F. Dean, A.P. Kowalczyk, and A.L. Mattheyses. 2022. Desmosomes undergo dynamic architectural changes during assembly and maturation. Tissue Barriers. 10. doi:10.1080/21688370.2021.2017225.

4. Belevich, I., M. Joensuu, D. Kumar, H. Vihinen, and E. Jokitalo. 2016. Microscopy Image Browser: A Platform for Segmentation and Analysis of Multidimensional Datasets. PLOS Biol. 14:e1002340. doi:10.1371/journal.pbio.1002340.

5. Bharathan, N.K., and A.J.G. Dickinson. 2019. Desmoplakin is required for epidermal integrity and morphogenesis in the Xenopus laevis embryo. Dev. Biol. 450:115–131. doi:10.1016/j.ydbio.2019.03.010.

6. Bharathan, N.K., W. Giang, C.L. Hoffman, J.S. Aaron, S. Khuon, T.L. Chew, S. Preibisch, E.T. Trautman, L. Heinrich, J. Bogovic, D. Bennett, D. Ackerman, W. Park, A. Petruncio, A.V. Weigel, S. Saalfeld, A. Wayne Vogl, S.N. Stahley, and A.P. Kowalczyk. 2023. Architecture and dynamics of a desmosome–endoplasmic reticulum complex. Nat. Cell Biol. 25:823–835. doi:10.1038/s41556-023-01154-4.

7. Bharathan, N.K., A.L. Mattheyses, and A.P. Kowalczyk. 2024. The desmosome comes into focus. J. Cell Biol. 223:e202404120. doi:10.1083/jcb.202404120.

8. Bizzozero, G. 1864. Delle cellule cigliate, del reticolo Malpighiano dell’ epidermide. Ann Univ Med. 190:110.

9. Bizzozero, G. 1870. Sulla struttura degli epiteli pavimentosi stratificati. Rendi Real Inst Lomb. 3:675.

10. Breathnach, A.S. 1975. Aspects Of Epidermal Ultrastructure. J. Invest. Dermatol. 65:2–15. doi:10.1111/1523-1747.ep12598018.

11. Breathnach, A.S., M. Gross, B. Martin, and C. Stolinski. 1976. A comparison of membrane fracture faces of fixed and unfixed glycerinated tissue. J. Cell Sci. 21:437–448. doi:10.1242/jcs.21.3.437.

12. Comet Technologies Canada Inc. (2025). Dragonfly 3D World. 2025.

13. Davis, J.D., and T.P. Wypych. 2021. Cellular and functional heterogeneity of the airway epithelium. Mucosal Immunol. 14:978–990. doi:10.1038/s41385-020-00370-7.

14. Devalia, J.L., R.J. Sapsford, C.W. Wells, P. Richman, and R.J. Davies. 1990. Culture and comparison of human bronchial and nasal epithelial cells in vitro. Respir. Med. 84:303–312. doi:10.1016/S0954-6111(08)80058-3.

15. Elias, P.M., and D.S. Friend. 1976. Vitamin-A-induced mucous metaplasia. An in vitro system for modulating tight and gap junction differentiation. J. Cell Biol. 68:173–188. doi:10.1083/jcb.68.2.173.

16. Fingerlin, T.E., E. Murphy, W. Zhang, A.L. Peljto, K.K. Brown, M.P. Steele, J.E. Loyd, G.P. Cosgrove, D. Lynch, S. Groshong, H.R. Collard, P.J. Wolters, W.Z. Bradford, K. Kossen, S.D. Seiwert, R.M. du Bois, C.K. Garcia, M.S. Devine, G. Gudmundsson, H.J. Isaksson, N. Kaminski, Y. Zhang, K.F. Gibson, L.H. Lancaster, J.D. Cogan, W.R. Mason, T.M. Maher, P.L. Molyneaux, A.U. Wells, M.F. Moffatt, M. Selman, A. Pardo, D.S. Kim, J.D. Crapo, B.J. Make, E.A. Regan, D.S. Walek, J.J. Daniel, Y. Kamatani, D. Zelenika, K. Smith, D. McKean, B.S. Pedersen, J. Talbert, R.N. Kidd, C.R. Markin, K.B. Beckman, M. Lathrop, M.I. Schwarz, and D.A. Schwartz. 2013. Genome-wide association study identifies multiple susceptibility loci for pulmonary fibrosis. Nat. Genet. 45:613–620. doi:10.1038/ng.2609.

17. Gloushankova, N.A., T. Wakatsuki, R.B. Troyanovsky, E. Elson, and S.M. Troyanovsky. 2003. Continual assembly of desmosomes within stable intercellular contacts of epithelial A-431 cells. Cell Tissue Res. 314:399–410. doi:10.1007/s00441-003-0812-3.

18. Godsel, L.M., S.N. Hsieh, E.V. Amargo, A.E. Bass, L.T. Pascoe-Mcgillicuddy, A.C. Huen, M.E. Thorne, C.A. Gaudry, J.K. Park, K. Myung, R.D. Goldman, T.-L. Chew, and K.J. Green. 2005. Desmoplakin assembly dynamics in four dimensions. J. Cell Biol. 171:1045–1059. doi:10.1083/jcb.200510038.

19. Hao, Y., S. Bates, H. Mou, J.H. Yun, B. Pham, J. Liu, W. Qiu, F. Guo, J.D. Morrow, C.P. Hersh, C.J. Benway, L. Gong, Y. Zhang, I.O. Rosas, M.H. Cho, J.-A. Park, P.J. Castaldi, F. Du, and X. Zhou. 2020. Genome-Wide Association Study: Functional Variant rs2076295 Regulates Desmoplakin Expression in Airway Epithelial Cells. Am. J. Respir. Crit. Care Med. 202:1225–1236. doi:10.1164/rccm.201910-1958OC.

20. He, W., P. Cowin, and D.L. Stokes. 2003. Untangling desmosomal knots with electron tomography. Science. 302:109–113. doi:10.1126/SCIENCE.1086957/SUPPL_FILE/HE.SOM.PDF.

21. Hegazy, M., A.L. Perl, S.A. Svoboda, and K.J. Green. 2022. Desmosomal Cadherins in Health and Disease. Annu. Rev. Pathol. Mech. Dis. 17:47–72. doi:10.1146/annurev-pathol-042320-092912.

22. Hunter, J.D. 2007. Matplotlib: A 2D graphics environment. Comput. Sci. Eng. 9:90–95. doi:10.1109/MCSE.2007.55.

23. Jorgens, D.M., J.L. Inman, M. Wojcik, C. Robertson, H. Palsdottir, W.T. Tsai, H. Huang, A. Bruni-Cardoso, C.S. López, M.J. Bissell, K. Xu, and M. Auer. 2017. Deep nuclear invaginations are linked to cytoskeletal filaments - integrated bioimaging of epithelial cells in 3D culture. J. Cell Sci. 130:177– 189. doi:10.1242/jcs.190967.

24. Kelly, D.E., and F.L. Shienvold. 1976. The desmosome: Fine structural studies with freeze-fracture replication and tannic acid staining of sectioned epidermis. Cell Tissue Res. 172:309–323. doi:10.1007/BF00399514/METRICS.

25. Lewis, J.D., A.L. Caldara, S.E. Zimmer, S.N. Stahley, A. Seybold, N.L. Strong, A.S. Frangakis, I. Levental, J.K. Wahl, A.L. Mattheyses, T. Sasaki, K. Nakabayashi, K. Hata, Y. Matsubara, A. Ishida-Yamamoto, M. Amagai, A. Kubo, and A.P. Kowalczyk. 2019. The desmosome is a mesoscale lipid raft–like membrane domain. Mol. Biol. Cell. 30:1390–1405. doi:10.1091/MBC.E18-10-0649/ASSET/IMAGES/LARGE/MBC-30-1390-G009.JPEG.

26. Lhamo, S., N.K. Bharathan, W. Giang, K.R. Levental, C.L. Simpson, S. Zimmer, and A.P. Kowalczyk. 2025. Cadherin Regulation of Endoplasmic Reticulum-Plasma Membrane Contact Sites. biorXiv 2025.10.15.682666. doi:10.1101/2025.10.15.682666.

27. Liu, Z., Q.P.H. Nguyen, Q. Guan, A. Albulescu, L. Erdman, Y. Mahdaviyeh, J. Kang, H. Ouyang, R.G. Hegele, T. Moraes, A. Goldenberg, S.D. Dell, and V. Mennella. 2020. A quantitative super-resolution imaging toolbox for diagnosis of motile ciliopathies. Sci. Transl. Med. 12:eaay0071. doi:10.1126/scitranslmed.aay0071.

28. Mathai, S.K., B.S. Pedersen, K. Smith, P. Russell, M.I. Schwarz, K.K. Brown, M.P. Steele, J.E. Loyd, J.D. Crapo, E.K. Silverman, D. Nickerson, T.E. Fingerlin, I.V. Yang, and D.A. Schwartz. 2016. Desmoplakin Variants Are Associated with Idiopathic Pulmonary Fibrosis. Am. J. Respir. Crit. Care Med. 193:1151–1160. doi:10.1164/rccm.201509-1863OC.

29. McDonald, K.L., and M. Auer. 2006. High-Pressure Freezing, Cellular Tomography, and Structural Cell Biology. BioTechniques. 41:137–143. doi:10.2144/000112226.

30. McDougall, C.M., M.G. Blaylock, J.G. Douglas, R.J. Brooker, P.J. Helms, and G.M. Walsh. 2008. Nasal Epithelial Cells as Surrogates for Bronchial Epithelial Cells in Airway Inflammation Studies. Am. J. Respir. Cell Mol. Biol. 39:560–568. doi:10.1165/rcmb.2007-0325OC.

31. Moch, M., N. Schwarz, R. Windoffer, and R.E. Leube. 2020. The keratin–desmosome scaffold: pivotal role of desmosomes for keratin network morphogenesis. Cell. Mol. Life Sci. CMLS. 77:543. doi:10.1007/S00018-019-03198-Y.

32. North, A.J., W.G. Bardsley, J. Hyam, E.A. Bornslaeger, H.C. Cordingley, B. Trinnaman, M. Hatzfeld, K.J. Green, A.I. Magee, and D.R. Garrod. 1999. Molecular map of the desmosomal plaque. J. Cell Sci. 112:4325–4336. doi:10.1242/jcs.112.23.4325.

33. Odland, G.F. 1958. The fine structure of the interrelationship of cells in the human epidermis. J. Biophys. Biochem. Cytol. 4:529–538. doi:10.1083/jcb.4.5.529.

34. Overton, J. 1962. Desmosome development in normal and reassociating cells in the early chick blastoderm. Dev. Biol. 4:532–548. doi:10.1016/0012-1606(62)90056-8.

35. Parslow, A., A. Cardona, and R.J. Bryson-Richardson. 2014. Sample Drift Correction Following 4D Confocal Time-lapse Imaging. J. Vis. Exp. JoVE. 51086. doi:10.3791/51086.

36. Perl, A.L., J.L. Pokorny, and K.J. Green. 2024. Desmosomes at a glance. J. Cell Sci. 137:jcs261899. doi:10.1242/jcs.261899.

37. Porter, K. 1956. Observations on the submicroscopic structure of animal epidermis. Proc. Int. Conf. Electron Microsc. 539.

38. Selby, C.C. 1955. An Electron Microscope Study Of The Epidermis Of Mammalian Skin In Thin Sections : I. Dermo-Epidermal Junction And Basal Cell Layer. J. Biophys. Biochem. Cytol. 1:429. doi:10.1083/JCB.1.5.429.

39. Selby, C.C. 1957. An electron microscope study of thin sections of human skin. II. Superficial cell layers of footpad epidermis. J. Invest. Dermatol. 29:131–149. doi:10.1038/JID.1957.80.

40. Sikora, M., U.H. Ermel, A. Seybold, M. Kunz, G. Calloni, J. Reitz, R. Martin Vabulas, G. Hummer, and A.S. Frangakis. 2020. Desmosome architecture derived from molecular dynamics simulations and cryo-electron tomography. Proc. Natl. Acad. Sci. U. S. A. 117:27132–27140. doi:10.1073/PNAS.2004563117/SUPPL_FILE/PNAS.2004563117.SM05.MOV.

41. Skepper, J.N. 1989. Membrane segregation in atrioventricular nodal myocytes of the golden hamster (Mesocricetus auratus). A cytochemical study using filipin and tomatine. J. Anat. 163:143–154.

42. Stahley, S.N., E.I. Bartle, C.E. Atkinson, A.P. Kowalczyk, and A.L. Mattheyses. 2016. Molecular organization of the desmosome as revealed by direct stochastic optical reconstruction microscopy. J. Cell Sci. 129:2897–2904. doi:10.1242/jcs.185785.

43. Stahley, S.N., and A.P. Kowalczyk. 2015. Desmosomes in acquired disease. Cell Tissue Res. 360:439–456. doi:10.1007/s00441-015-2155-2.

44. Stahley, S.N., M. Saito, V. Faundez, M. Koval, A.L. Mattheyses, and A.P. Kowalczyk. 2014. Desmosome assembly and disassembly are membrane raft-dependent. PloS One. 9. doi:10.1371/JOURNAL.PONE.0087809.

45. Vanslembrouck, B., A. Kremer, B. Pavie, F. van Roy, S. Lippens, and J. van Hengel. 2018. Three-dimensional reconstruction of the intercalated disc including the intercellular junctions by applying volume scanning electron microscopy. Histochem. Cell Biol. 149:479–490. doi:10.1007/S00418-018-1657-X/TABLES/2.

46. Vanslembrouck, B., A. Kremer, F. van Roy, S. Lippens, and J. van Hengel. 2020. Unravelling the ultrastructural details of αT-catenin-deficient cell-cell contacts between heart muscle cells by the use of FIB-SEM. J. Microsc. 279:189–196. doi:10.1111/JMI.12855.

47. Vijayakumaran, A., C. Godbehere, A. Abuammar, S.Y. Breusegem, L.R. Hurst, N. Morone, J. Llodra, M.T. Dalbay, N.M. Tanvir, K. MacLellan-Gibson, C. O’Callaghan, E. Lorentzen, C.P. Team, F.-S. Technology, A.J. Murray, K. Narayan, and V. Mennella. 2024. 3D nanoscale architecture of the respiratory epithelium reveals motile cilia-rootlets-mitochondria axis of communication. biorXiv 2024.09.08.611854. doi:10.1101/2024.09.08.611854.

48. Waskom, M. 2021. seaborn: statistical data visualization. J. Open Source Softw. 6:3021. doi:10.21105/joss.03021.

49. Windoffer, R., M. Borchert-Stuhlträger, and R.E. Leube. 2002. Desmosomes: interconnected calcium-dependent structures of remarkable stability with significant integral membrane protein turnover. J. Cell Sci. 115:1717–1732. doi:10.1242/JCS.115.8.1717.

50. Wolburg, H., R. Kästner, and G. Kurz-Isler. 1983. Lack of orthogonal particle assemblies and presence of tight junctions in astrocytes of the goldfish (Carassius auratus). Cell Tissue Res. 234:389–402. doi:10.1007/BF00213776.

51. Xu, C.S., K.J. Hayworth, Z. Lu, P. Grob, A.M. Hassan, J.G. García-Cerdán, K.K. Niyogi, E. Nogales, R.J. Weinberg, and H.F. Hess. 2017. Enhanced FIB-SEM systems for large-volume 3D imaging. eLife. 6. doi:10.7554/elife.25916.

52. Zimmer, S.E., and A.P. Kowalczyk. 2020. The desmosome as a model for lipid raft driven membrane domain organization. Biochim. Biophys. Acta BBA - Biomembr. 1862:183329. doi:10.1016/j.bbamem.2020.183329.

53. Zimmer, S.E., and A.P. Kowalczyk. 2024. The desmosome as a dynamic membrane domain. Curr. Opin. Cell Biol. 90:102403. doi:10.1016/j.ceb.2024.102403.

